# A human monoclonal antibody blocking SARS-CoV-2 infection

**DOI:** 10.1101/2020.03.11.987958

**Authors:** Chunyan Wang, Wentao Li, Dubravka Drabek, Nisreen M.A. Okba, Rien van Haperen, Albert D.M.E. Osterhaus, Frank J.M. van Kuppeveld, Bart L. Haagmans, Frank Grosveld, Berend-Jan Bosch

## Abstract

The emergence of the novel human coronavirus SARS-CoV-2 in Wuhan, China has caused a worldwide epidemic of respiratory disease (COVID-19). Vaccines and targeted therapeutics for treatment of this disease are currently lacking. Here we report a human monoclonal antibody that neutralizes SARS-CoV-2 (and SARS-CoV). This cross-neutralizing antibody targets a communal epitope on these viruses and offers potential for prevention and treatment of COVID-19.

## Main text

The severe acute respiratory syndrome coronavirus 2 (SARS-CoV-2) is the etiological agent of the coronavirus induced disease 19 (COVID-19) that emerged in China late 2019 and causing a worldwide epidemic^1^. As of March 10^th^ 2020, over 115,113 cases have been reported in 109 countries, of which 4,063 (3.5%) succumbed to the infection^2^. SARS-CoV-2 belongs to the *Sarbecovirus* subgenus (genus *Betacoronavirus*, family *Coronaviridae*)^3^ together with SARS-CoV that emerged in 2002 causing approximately 8000 infections with a lethality of 10%. Both viruses crossed species barriers from an animal reservoir and can cause a life-threatening respiratory illness in humans. Presently no approved targeted therapeutics are available for COVID-19. Monoclonal antibodies targeting vulnerable sites on viral surface proteins are increasingly recognised as a promising class of drugs against infectious diseases and have shown therapeutic efficacy for a number of viruses^4, 5^.

Coronavirus neutralizing antibodies primarily target the trimeric spike (S) glycoproteins on the viral surface that mediate entry into host cells. The S protein has two functional subunits that mediate cell attachment (the S1 subunit, existing of four core domains S1_A_ through S1_D_) and fusion of the viral and cellular membrane (the S2 subunit). Potent neutralizing antibodies often target the receptor interaction site in S1, disabling receptor interactions^6–11^. The spike proteins of SARS-CoV-2 (SARS2-S; 1,273 residues, strain Wuhan-Hu-1) and SARS-CoV (SARS-S, 1,255 residues, strain Urbani) are 77.5% identical by primary amino acid sequence, are structurally very similar^12, 13^ and commonly bind the human angiotensin coverting enzyme 2 (ACE2) protein as a host receptor^1, 14^ through their S1_B_ domain. Receptor interaction is known to trigger irreversible conformational changes in coronavirus spike proteins enabling membrane fusion^15^.

In order to identify SARS-CoV-2 neutralizing antibodies, ELISA-(cross)reactivity was assessed of antibody-containing supernatants of a collection of 51 SARS-S hybridoma’s derived from immunized transgenic H2L2 mice that encode chimeric immunoglobulins with human variable heavy and light chains and constant regions of rat origin (Suppl.Fig.1). Four of 51 SARS-S hybridoma supernatants displayed ELISA-cross-reactivity with the SARS2-S1 subunit (S residues 1-681; Suppl.Fig.1), of which one (47D11) exhibited cross-neutralizing activity of SARS-S and SARS2-S pseudotyped VSV infection. The chimeric 47D11 H2L2 antibody was reformatted and recombinantly expressed as a fully human IgG1 isotype antibody for further characterization.

The human 47D11 antibody binds to cells expressing the full-length spike proteins of SARS-CoV and SARS-CoV-2 (Fig.1a). The 47D11 antibody was found to potently inhibit infection of VeroE6 cells with SARS-S and SARS2-S pseudotyped VSV with IC_50_ values of 0.06 and 0.08 μg/ml (Fig.1b), respectively. Authentic infection of VeroE6 cells with SARS-CoV and SARS-CoV-2 was neutralized with IC_50_ values of 0.19 and 0.57 μg/ml (Fig.1c). Using ELISA 47D11 was shown to target the S1_B_ receptor binding domain (RBD) of SARS-S and SARS2-S. 47D11 bound the S1_B_ of both viruses with similar affinities as shown by the ELISA-based half maximal effective concentration (EC_50_) values (0.02 and 0.03 μg/ml, respectively; Fig.2a). ELISA-based binding affinity of 47D11 for the spike ectodomain (S_ecto_) of SARS-CoV was higher relative to that of SARS-CoV-2 (EC_50_ values: 0.018 and 0.15 μg/ml, respectively), despite equimolar antigen coating (Suppl.Fig.2). Congruent with the ELISA-reactivities, measurement of binding kinetics of 47D11 by biolayer interferometry showed that 47D11 binds SARS-S_ecto_ with higher affinity (equilibrium dissociation constant [*K_D_*]: 0.745 nM) relative to SARS2-S_ecto_ (*K_D_* 10.8 nM) whereas affinity for SARS-S1_B_ and SARS2-S1_B_ was in a similar range (16.1 and 9.6 nM, respectively, Suppl.Fig.3). This difference may originate from differences in epitope accessibility in SARS-S versus SARS2-S, as domain B can adopt a closed and open conformation in the prefusion spike homotrimer^12, 13^. Remarkably, binding of 47D11 to SARS-S1_B_ and SARS2-S1_B_ did not compete with S1_B_ binding to the ACE2 receptor expressed at the cell surface as shown by flow cytometry (Fig.2b; Suppl.Fig.4) nor with S_ecto_ and S1_B_ binding to soluble ACE2 in solid-phase based assay (Suppl.Fig.5), whereas two SARS-S1 specific antibodies 35F4 and 43C6 that neutralize SARS-S (but not SARS2-S) pseudotyped VSV infection (Suppl.Fig.6) do block binding of SARS-S_ecto_ and SARS-S1_B_ to ACE2. Using a trypsin-triggered cell-cell fusion assay, 47D11 was shown to impair SARS-S and SARS2-S mediated syncytia formation (Suppl.Fig.7). Our data show that 47D11 neutralizes SARS-CoV and SARS-CoV-2 through a yet unknown mechanism that is different from receptor binding interference. Alternative mechanisms of coronavirus neutralization by RBD-targeting antibodies have been reported including spike inactivation through antibody-induced destabilization of its prefusion structure^15^, which may also apply for 47D11.

**Fig.1.**
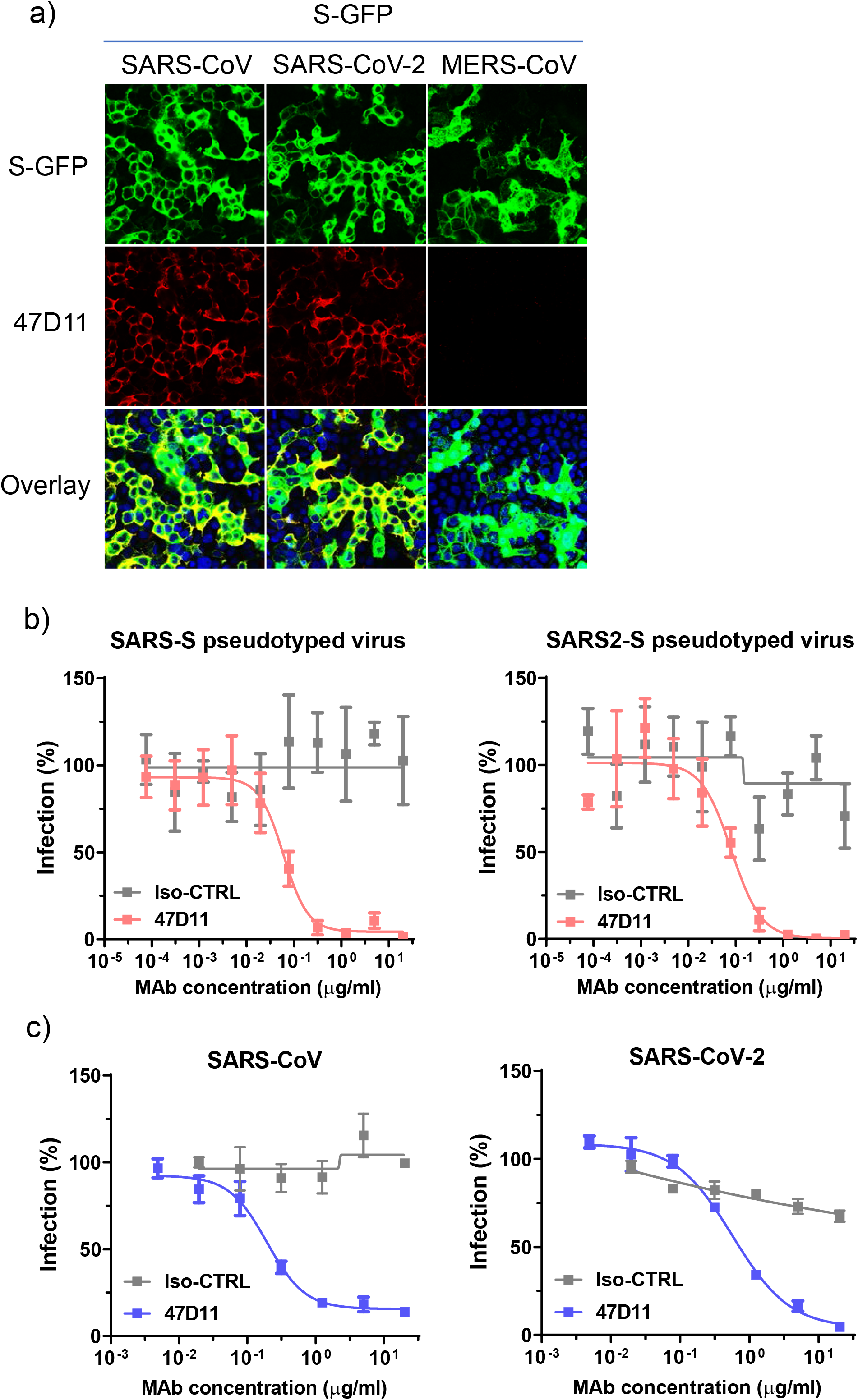
47D11 neutralizes SARS-CoV and SARS-CoV-2. a) Binding of 47D11 to HEK-293T cells expressing GFP-tagged spike proteins of SARS-CoV and SARS-CoV-2 detected by immunofluorescence assay. The human mAb 7.7G6 targeting the MERS-CoV S1_B_ spike domain was taken along as a negative control, cell nuclei in the overlay images are visualized with DAPI. b) Antibody-mediated neutralization of infection of luciferase-encoding VSV particles pseudotyped with spike proteins of SARS-CoV and SARS-CoV-2. Pseudotyped VSV particles pre-incubated with antibodies at indicated concentrations (see methods) were used to infect VeroE6 cells and luciferase activities in cell lysates were determined at 24 h post transduction to calculate infection (%) relative to non-antibody-treated controls. The average ± SD from at least two independent experiments performed is shown. Iso-CTRL: irrelevant isotype monoclonal antibody. c) Antibody-mediated neutralization of SARS-CoV and SARS-CoV-2 infection on VeroE6 cells. The experiment was performed with triplicate samples, the average ± SD is shown.

**Fig.2.**
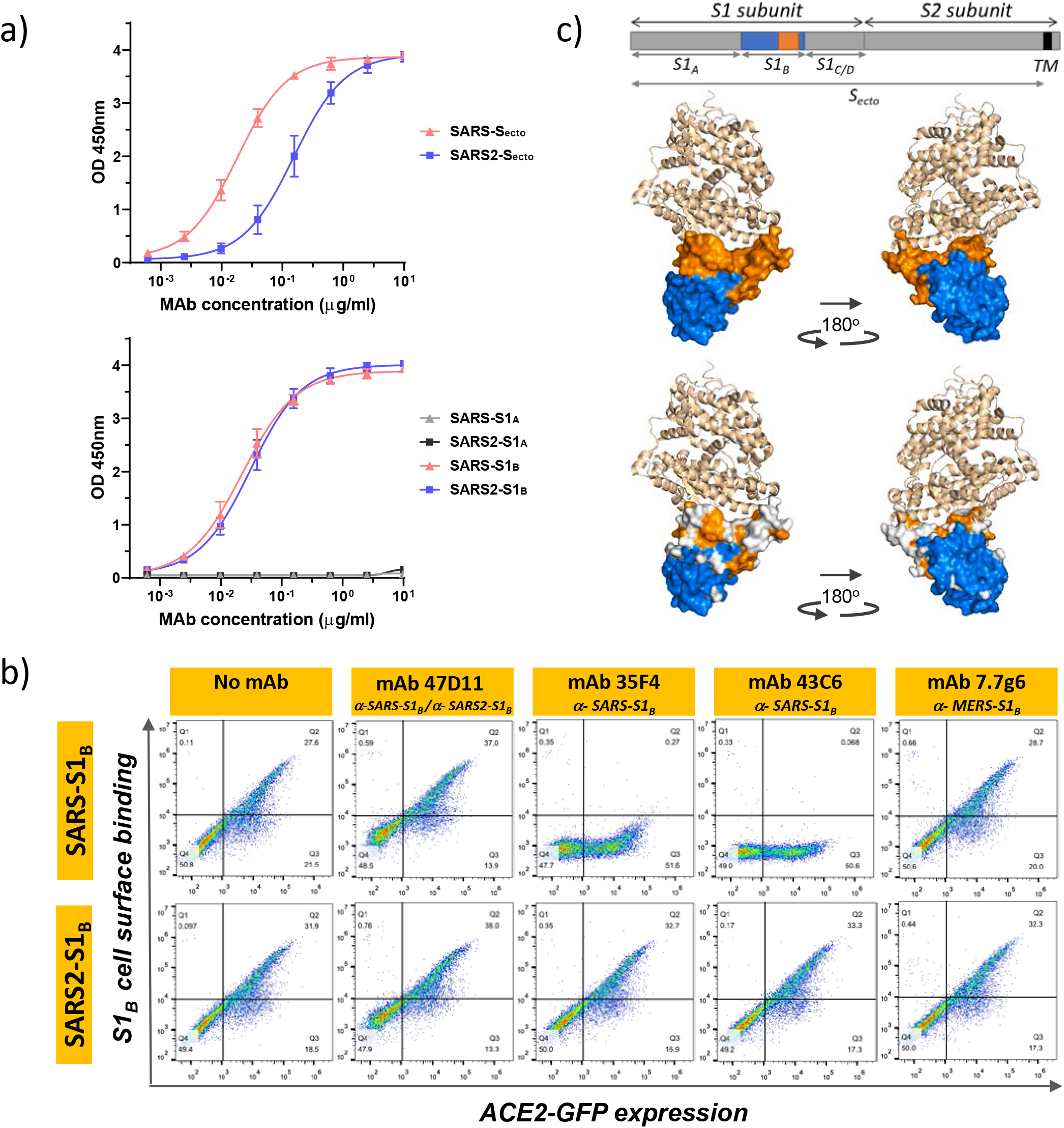
The neutralizing 47D11 monoclonal antibody binds the receptor binding domain of SARS-CoV and SARS-CoV-2 spike proteins without eliminating S1_B_/ACE2 receptor interaction. a) ELISA binding curves of 47D11 to S_ecto_ (upper panel) or S1_A_ and S1_B_ (RBD) (lower panel) of SARS-S and SARS2-S coated at equimolar concentrations. The average ± SD from at least two independent experiments performed is shown. b) Interference of antibodies with binding of the S-S1_B_ of SARS-CoV and SARS-CoV-2 to cell surface ACE2-GFP analysed by flow cytometry. Prior to cell binding, S1_B_ was mixed with mAb (mAbs 47D11, 35F4, 43C6, 7.7G6, in H2L2 format) with indicated specificity in a mAb:S1_B_ molar ratio of 8:1 (see Suppl.Fig.4 for an extensive analysis using different mAb:S1_B_ molar ratio’s). Cells are analysed for (ACE2-)GFP expression (x-axis) and S1_B_ binding (y-axis). Percentages of cells that scored negative, single positive, or double positive are shown in each quadrant. c) Divergence in surface residues in S1_B_ of SARS-CoV and SARS-CoV-2. Upper panel: Structure of the SARS-CoV spike protein S1_B_ RBD in complex with human ACE2 receptor (PDB: 2AJF)^18^. ACE2 (wheat color) is visualized in ribbon presentation. The S1_B_ core domain (blue) and subdomain (orange) are displayed in surface presentation using PyMOL, and are visualized with the same colors in the linear diagram of the spike protein above, with positions of the S1 and S2 subunits, the S ectodomain (S_ecto_), the S1 domains S1_A-D_ and the transmembrane domain (TM) indicated. Lower panel: Similar as panel above with surface residues on S1_B_ of SARS-CoV that are at variance with SARS-CoV-2 colorored in white.

The SARS2-S1_B_ receptor binding domain (residues 338-506) consists of a core domain and a receptor binding subdomain (residues 438-498) looping out from the antiparallel betasheet core domain structure that directly engages the receptor. Compared to the S1_B_ core domain, the protein sequence identity of the S1_B_ receptor interacting subdomain of SARS-S and SARS2-S is substantially lower (46.7% versus 86.3%; Suppl.Fig.8 and Fig.2c). Potent neutralizing antibodies often target this receptor binding subdomain. However, due to common variations in this subdomain, these antibodies are often virus-specific and bind and neutralize related viruses poorly^16, 17^. The cross-reactive nature of 47D11 indicates that the antibody is more likely to target the conserved core structure of the S1_B_ receptor binding domain. S1_B_ binding by 47D11 further away from the receptor binding interface explains its inability to compromise spike-receptor interaction.

In conclusion, this is the first report on a (human) monoclonal antibody that neutralizes SARS-CoV-2. 47D11 binds a conserved epitope on the spike receptor binding domain explaining its ability to cross-neutralize SARS-CoV and SARS-CoV-2, using a mechanism that is independent of receptor binding inhibition. This antibody will be useful for development of antigen detection tests and serological assays targeting SARS-CoV-2. Neutralizing antibodies can alter the course of infection in the infected host supporting virus clearance or protect an uninfected host that is exposed to the virus^4^. Hence, this antibody offers the potential to prevent and/or treat COVID-19, and possibly also other future emerging diseases in humans caused by viruses from the *Sarbecovirus* subgenus.

## Acknowledgements

We thank dr. Yoshiharu Matsuura (Osaka University, Japan) for the provision of the luciferase-encoding VSV-G pseudotyped VSVΔG-luc virus, and Yongle Yang, Michael van der Reijden and Rick Janssens for technical support. We thank Christian Drosten (Charité Universitätsmedizin Berlin, Germany) for provision of the SARS-CoV-2 virus. This study was done within the framework of National Centre for One Health (NCOH) and the Innovative Medicines Initiative (IMI) Zoonotic Anticipation and Preparedness Initiative [ZAPI project; grant agreement no. 115760]. The mice used in this study were provided by Harbour Antibodies BV, a daughter company of Harbour Biomed (http://www.harbourbiomed.com). C. Wang was supported by a grant from the Chinese Scholarship Council (file number CSC201708620178).

## Author Contributions

B.J.B. designed and coordinated the study. C.W., W.L., N.M.A.O., R.v.H. and D.D conducted the experiments. D.D., B.L.H. and B.J.B. supervised part of the experiments. All authors contributed to the interpretations and conclusions presented. B.J.B. wrote the manuscript, B.L.H., F.J.M.K., A.D.M.E.O. and F.G. participated in editing the manuscript.

## Materials and Methods

### Expression and purification of coronavirus spike proteins

Coronavirus spike ectodomains (S_ecto_) of SARS-CoV (residues 1–1,182; strain CUHK-W1; GenBank: AAP13567.1) and SARS-CoV-2 (residues 1–1,213; strain Wuhan-Hu-1; GenBank: QHD43416.1) were expressed transiently in HEK-293T cells with a C-terminal trimerization motif and Strep-tag using the pCAGGS expression plasmid. Similarly, pCAGGS expression vectors encoding S1 or its subdomains of SARS-CoV (S1, residues 1-676; S1_A_, residues 1-302; S1_B_, residues, 325-533), and SARS-CoV-2 (S1, residues 1-682; S1_A_, residues 1-294; S1_B_, residues 329-538) C-terminally tagged with Fc domain of human or mouse IgG or Strep-tag were generated as described before^19^. Recombinant proteins were expressed transiently in HEK-293T cells and affinity purified from the culture supernatant by protein-A sepharose beads (GE Healthcare) or streptactin beads (IBA) purification. Purity and integrity of all purified recombinant proteins was checked by coomassie stained SDS-PAGE.

### Generation of H2L2 mAbs

H2L2 mice were sequentially immunized in two weeks intervals with purified S_ecto_ of different CoVs in the following order: HCoV-OC43, SARS-CoV, MERS-CoV, HCoV-OC43, SARS-CoV and MERS-CoV. Antigens were injected at 20-25 μg/mouse using Stimune Adjuvant (Prionics) freshly prepared according to the manufacturer instruction for first injection, while boosting was done using Ribi (Sigma) adjuvant. Injections were done subcutaneously into the left and right groin each (50 μl) and 100 μl intraperitoneally. Four days after the last injection, spleen and lymph nodes are harvested, and hybridomas made by standard method using SP 2/0 myeloma cell line (ATCC#CRL-1581) as a fusion partner. Hybridomas were screened in antigen-specific ELISA and those selected for further development, subcloned and produced on a small scale (100 ml of medium). For this purpose, hybridomas are cultured in serum- and protein-free medium for hybridoma culturing (PFHM-II (1X), Gibco) with addition of non-essential amino acids 100X NEAA, Biowhittaker Lonza, Cat BE13-114E). H2L2 antibodies were purified from hybridoma culture supernatants using Protein-A affinity chromatography. Purified antibodies were stored at 4°C until use.

### Production of human monoclonal antibody 47D11

For recombinant human mAb production, the cDNA’s encoding the 47D11 H2L2 mAb variable regions of the heavy and light chains were cloned into expression plasmids containing the human IgG1 heavy chain and Ig kappa light chain constant regions, respectively (InvivoGen). Both plasmids contain the interleukin-2 signal sequence to enable efficient secretion of recombinant antibodies. Recombinant human 47D11 mAb and previously described Isotype-control (anti-Streptag mAb) or 7.7G6 mAb were produced in HEK-293T cells following transfection with pairs of the IgG1 heavy and light chain expression plasmids according to protocols from InvivoGen. Human antibodies were purified from cell culture supernatants using Protein-A affinity chromatography. Purified antibodies were stored at 4°C until use.

### Immunofluorescence microscopy

Antibody binding to cell surface spike proteins of SARS-CoV, SARS-CoV-2 and MERS-CoV was measured by immunofluoresence microscopy. HEK-293T cells seeded on glass slides were transfected with plasmids encoding SARS-S, SARS2-S or MERS-S - C-terminally fused to the green fluorescence protein (GFP) - using Lipofectamine 2000 (Invitrogen). Two days post transfection, cells were fixed by incubation with 2% paraformaldehyde in PBS for 20 min at room temperature and stained for nuclei with 4,6-diamidino-2-phenylindole (DAPI). Cells were subsequently incubated with mAbs at a concentration of 10 μg/ml for 1 h at room temperature, followed by incubation with Alexa Fluor 594 conjugated goat anti-human IgG antibodies (Invitrogen, Thermo Fisher Scientific) for 45 min at room temperature. The fluorescence images were recorded using a Leica SpeII confocal microscope.

### Flow cytometry-based receptor binding inhibition assay

Antibody interference of S1_B_ binding to human ACE2 receptor on the cell surface was measured by flow cytometry. HEK-293T cells were seeded at a density of 2.5×10^5^ cells per ml in a T75 flask. After reaching 70~80% confluency, cells were transfected with an expression plasmid encoding human ACE2 - C-terminally fused to the GFP - using Lipofectamine 2000 (Invitrogen). Two days post transfection, cells were dissociated by cell dissociation solution (Sigma-aldrich, Merck KGaA; cat. no. C5914). 2.5 μg/ml of human Fc tagged SARS-S1_B_ and SARS2-S1_B_ was preincubated with mAb at the indicated mAb:S1_B_ molar ratios for 1 hour on ice and subjected to flow cytometry. Single cell suspensions in FACS buffer were centrifuged at 400×g for 10 min. Cells were subsequently incubated with S1_B_ and mAb mixture for 1 h on ice, followed by incubation with Alexa Fluor 594 conjugated goat anti-human IgG antibodies (Invitrogen, Thermo Fisher Scientific) for 45 min at room temperature. Cells were subjected to flow cytometric analysis with a CytoFLEX Flow Cytometer (Beckman Coulter). The results were analysed by FlowJo (version 10).

### Pseudotyped virus neutralization assay

Production of VSV pseudotyped with SARS-S and SARS2-S was performed as described previously with some adaptations^11^. Briefly, HEK-293T cells were transfected with pCAGGS expression vectors encoding SARS-S or SARS2-S carrying a 28- or 18-a.a. cytoplasmic tail truncation, respectively. One day post transfection, cells were infected with the VSV-G pseudotyped VSVΔG bearing the firefly (*Photinus pyralis*) luciferase reporter gene. Twenty-four hours later, supernatants containing SARS-S/SARS2-S pseudotyped VSV particles were harvested and titrated on African green monkey kidney VeroE6 cells. In the virus neutralization assay, mAbs were serially diluted at two times the desired final concentration in DMEM supplemented with 1% fetal calf serum (Bodinco), 100 U/ml Penicillin and 100 μg/ml Streptomycin. Diluted mAbs were incubated with an equal volume of pseudotyped VSV particles for 1 hour at room temperature, inoculated on confluent VeroE6 monolayers in 96-well plated, and further incubated at 37°C for 24 hours. Luciferase activity was measured on a Berthold Centro LB 960 plate luminometer using D-luciferin as a substrate (Promega). The percentage of infectivity was calculated as ratio of luciferase readout in the presence of mAbs normalized to luciferase readout in the absence of mAb. The half maximal inhibitory concentrations (IC_50_) were determined using 4-parameter logistic regression (GraphPad Prism version 8).

### Virus neutralization assay

Neutralization of authentic SARS-CoV and SARS-CoV-2 was performed using a plaque reduction neutralization test (PRNT) as described earlier, with some modifications^20^. In brief, mAbs were two-fold serially diluted and mixed with SARS-CoV or SARS-CoV-2 for 1 hour. The mixture was then added to VeroE6 cells and incubated for 1 hr, after which the cells were washed and further incubated in medium for 8 hrs. The cells were then fixed and stained using a rabbit anti-SARS-CoV serum (Sino Biological) and a secondary peroxidase-labelled goat anti-rabbit IgG (Dako). The signal was developed using a precipitate forming TMB substrate (True Blue, KPL) and the number of infected cells per well were counted using the ImmunoSpot® Image analyzer (CTL Europe GmbH). The half maximal inhibitory concentrations (IC_50_) were determined using 4-parameter logistic regression (GraphPad Prism version 8).

### ELISA analysis of antibody binding to CoV spike antigens

NUNC Maxisorp plates (Thermo Scientific) were coated with equimolar antigen amounts at 4°C overnight. Plates were washed three times with Phosphate Saline Buffer (PBS) containing 0.05% Tween-20 and blocked with 3% Bovine Serum Albumin (BSA) in PBS containing 0.1% Tween-20 at room temperature for 2 hours. Four-folds serial dilutions of mAbs starting at 10 μg/ml (diluted in blocking buffer) were added and plates were incubated for 1 hour at room temperature. Plates were washed three times and incubated with HRP-conjugated goat anti-human secondary antibody (ITK Southern Biotech) diluted 1:2000 in blocking buffer for 1 hour at room temperature. An HRP-conjugated anti-StrepMAb (IBA, Cat.no: 2-1509-001) antibody was used to corroborate equimolar coating of the Strep-tagged spike antigens. HRP activity was measured at 450 nanometer using tetramethylbenzidine substrate (BioFX) and an ELISA plate reader (EL-808, Biotek). Half-maximum effective concentration (EC_50_) binding values were calculated by non-linear regression analysis on the binding curves using GraphPad Prism (version 8).

---

**Suppl. Fig.1.**
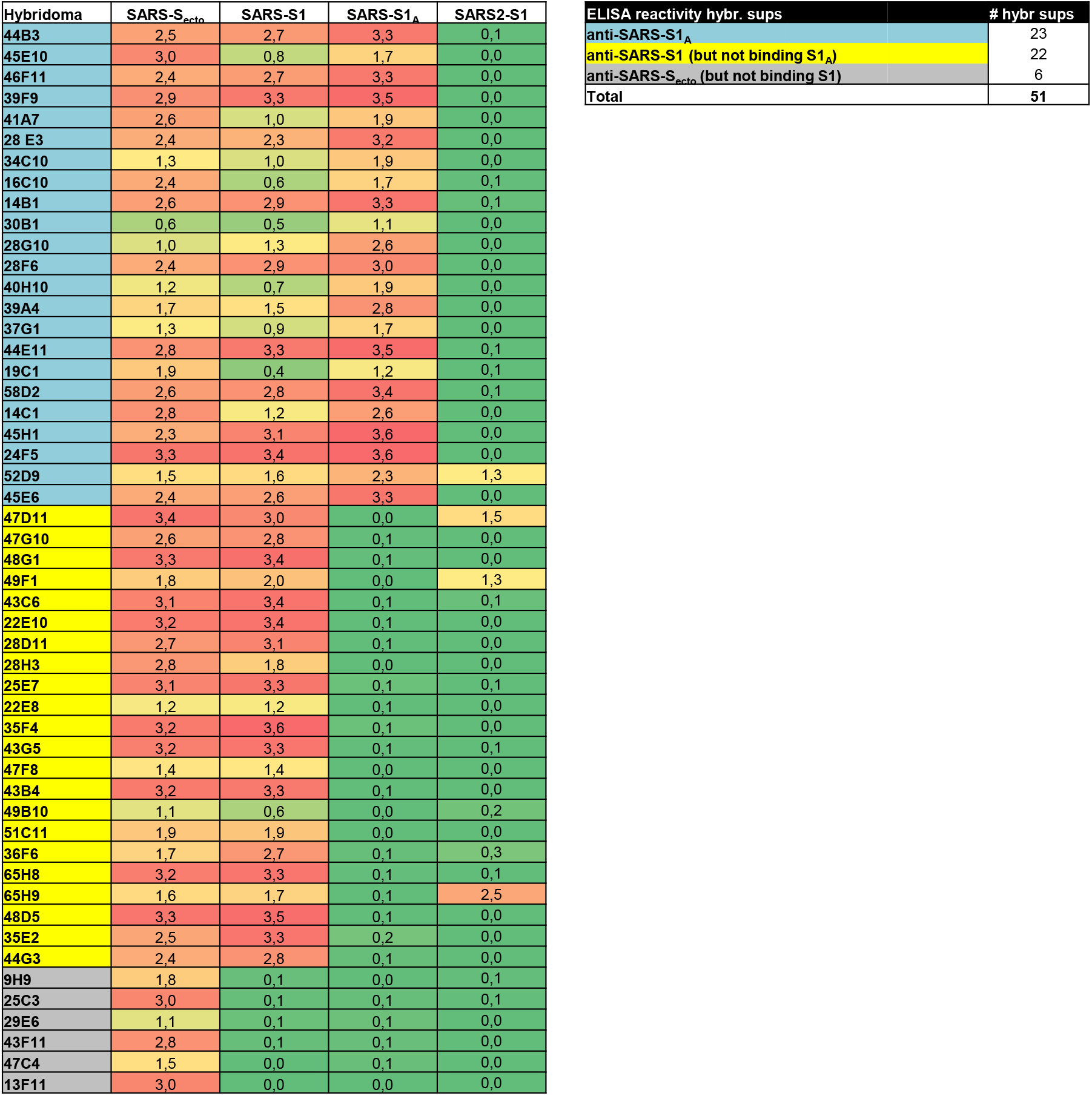
ELISA cross-reactivity of antibody-containing supernatants of SARS-S H2L2 hybridomas towards SARS2-S1. SARS-S targeting hybridomas developed by conventional hybridoma technology from immunized H2L2 transgenic mice (Harbour Biomed), as described before^1^. These mice - carrying encoding the heavy and light chain human immunoglobulin repertoire - were sequentially immunized with 2-week intervals with trimeric spike protein ectodomains (S_ecto_) of three human coronaviruses from the betacoronavirus genus in the following order: 1. HCoV-OC43-S_ecto_, 2. SARS-CoV-S_ecto_, 3. MERS-CoV-S_ecto_, 4. HCoV-OC43-S_ecto_, 5. SARS-CoV-S_ecto_, 6. MERS-CoV-S_ecto_. Four days after the last immunization, splenocytes and lymph node lymphocytes were harvested and hybridomas were generated. Antibodies in the cell supernatants were tested for ELISA-reactivity against SARS-S_ecto_, SARS-S1, SARS-S1_A_ and SARS2-S1. Of the 51 hybridoma supernatants that reacted with SARS-S_ecto_ only, 23 reacted with SARS-S1_A_, 22 with SARS-S1 but not SARS-S1_A_, 6 with SARS-S_ecto_ but not SARS-S1. Four of the 51 SARS-S_ecto_ hybridoma supernatants reacted with SARS2-S1 (see column on the right). The table displays ELISA-signal intensities (OD_450nm_ values) of hybridoma supernatants for the different antigens.

**Suppl. Fig.2.**
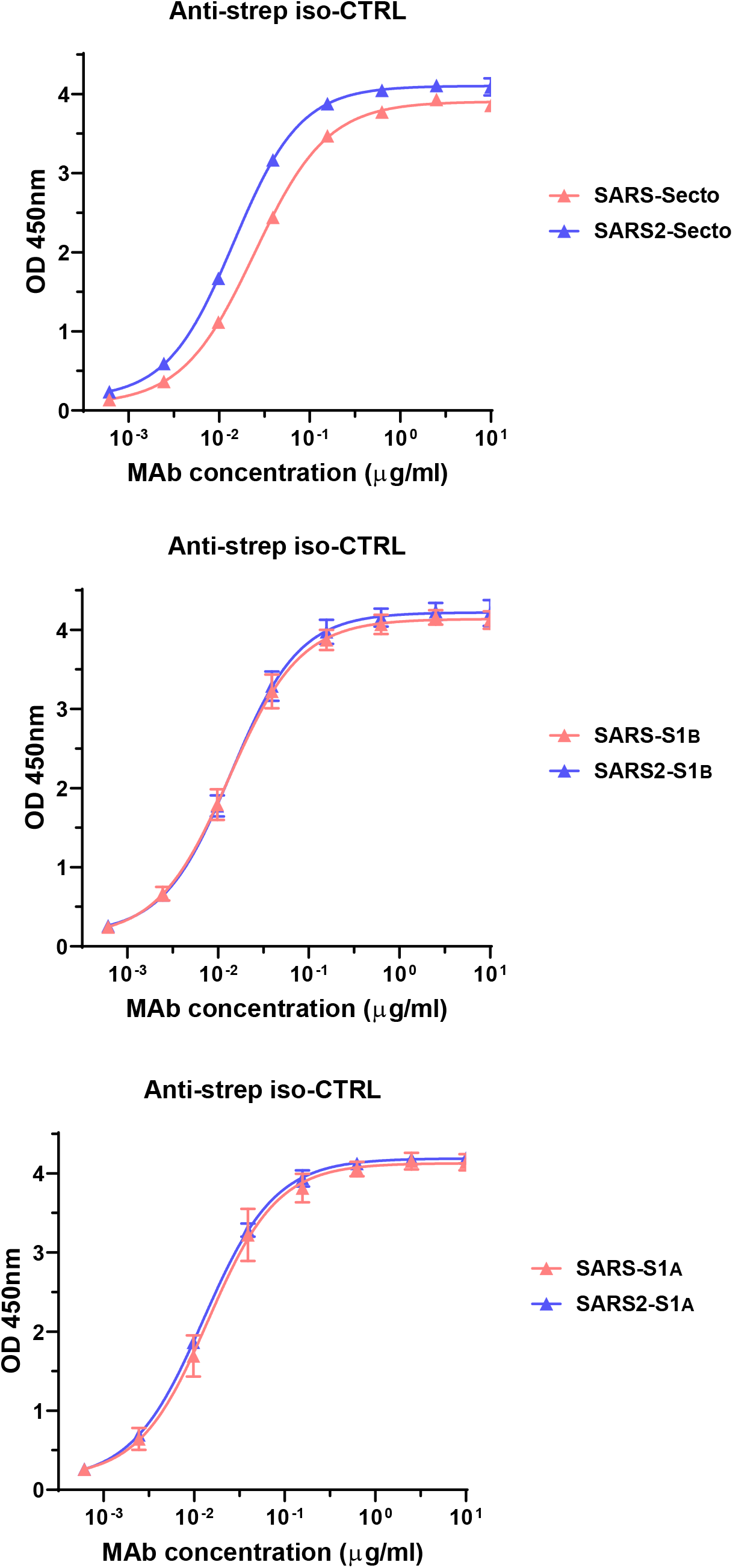
ELISA binding curve of the anti-StrepMAb (IBA) antibody to Strep-tagged spike antigens to corroborate equimolar ELISA plate coating of SARS-S_ecto_ / SARS2-S_ecto_ (upper panel), SARS-S1_B_ / SARS2-S1_B_ (middle panel) and SARS-S1_A_ / SARS2-S1_A_ (lower panel) antigens used in Fig.2a.

**Suppl. Fig.3.**
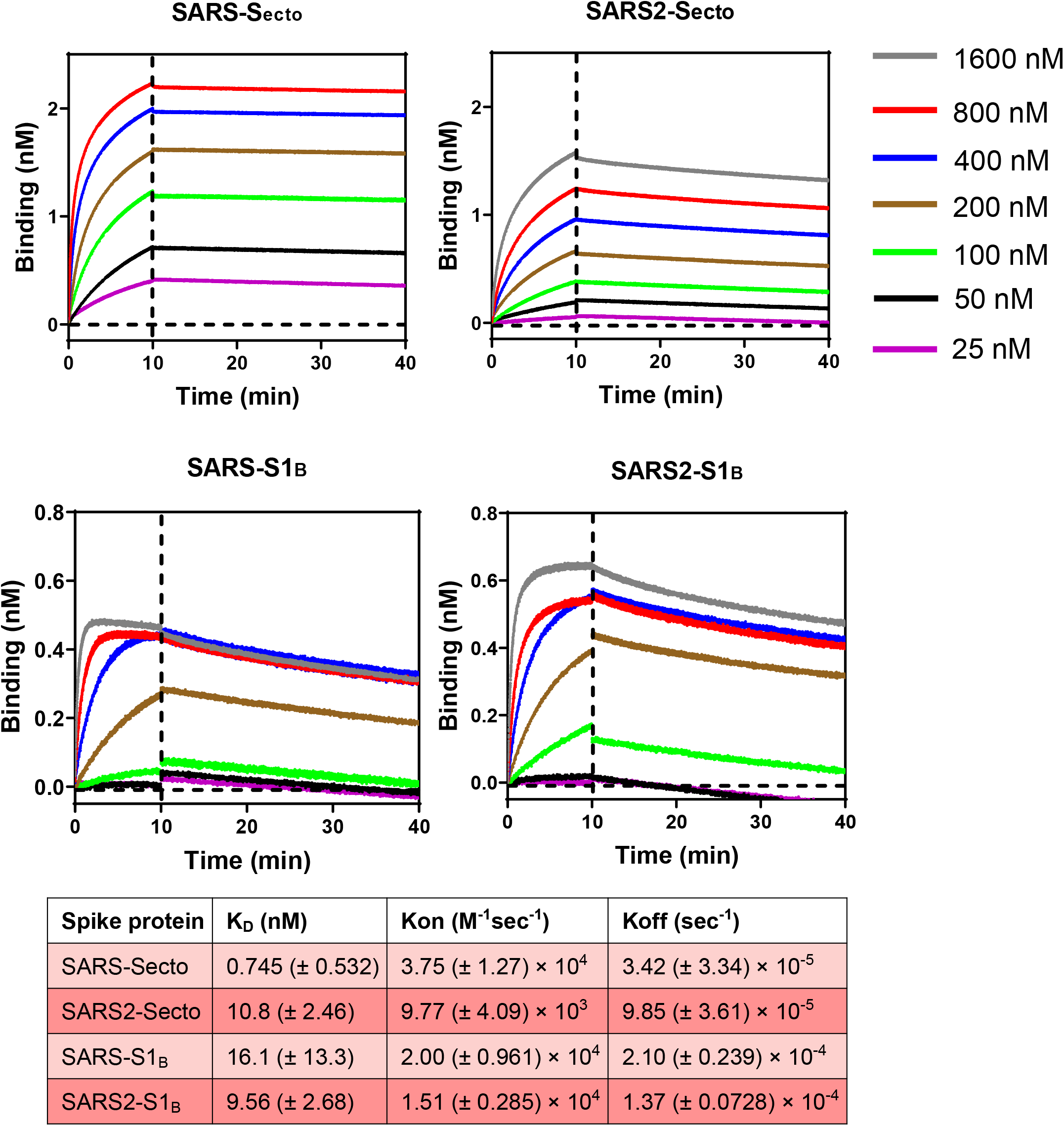
Binding kinetics of 47D11 to the S ectodomain and S1_B_ of SARS-CoV and SARS-CoV-2. Binding kinetics of 47D11 to immobilized recombinant SARS-S_ecto_, SARS2-S_ecto_, SARS-S1_B_ and SARS2-S1_B_ was measured using biolayer interferometry at 25°C, as described previously^21^. Kinetic binding assay was performed by loading 47D11 mAb at optimal concentration (42 nM) on anti-human Fc biosensor for 10 mins. Antigen association step was performed by incubating the sensor with a range of concentrations of the recombinant spike ectodomain (1600-800-400-200-100-50-25 nM) for 10 min, followed by a dissociation step in PBS for 60 min. The kinetics constants were calculated using 1:1 Langmuir binding model on Fortebio Data Analysis 7.0 software.

**Suppl. Fig.4.**
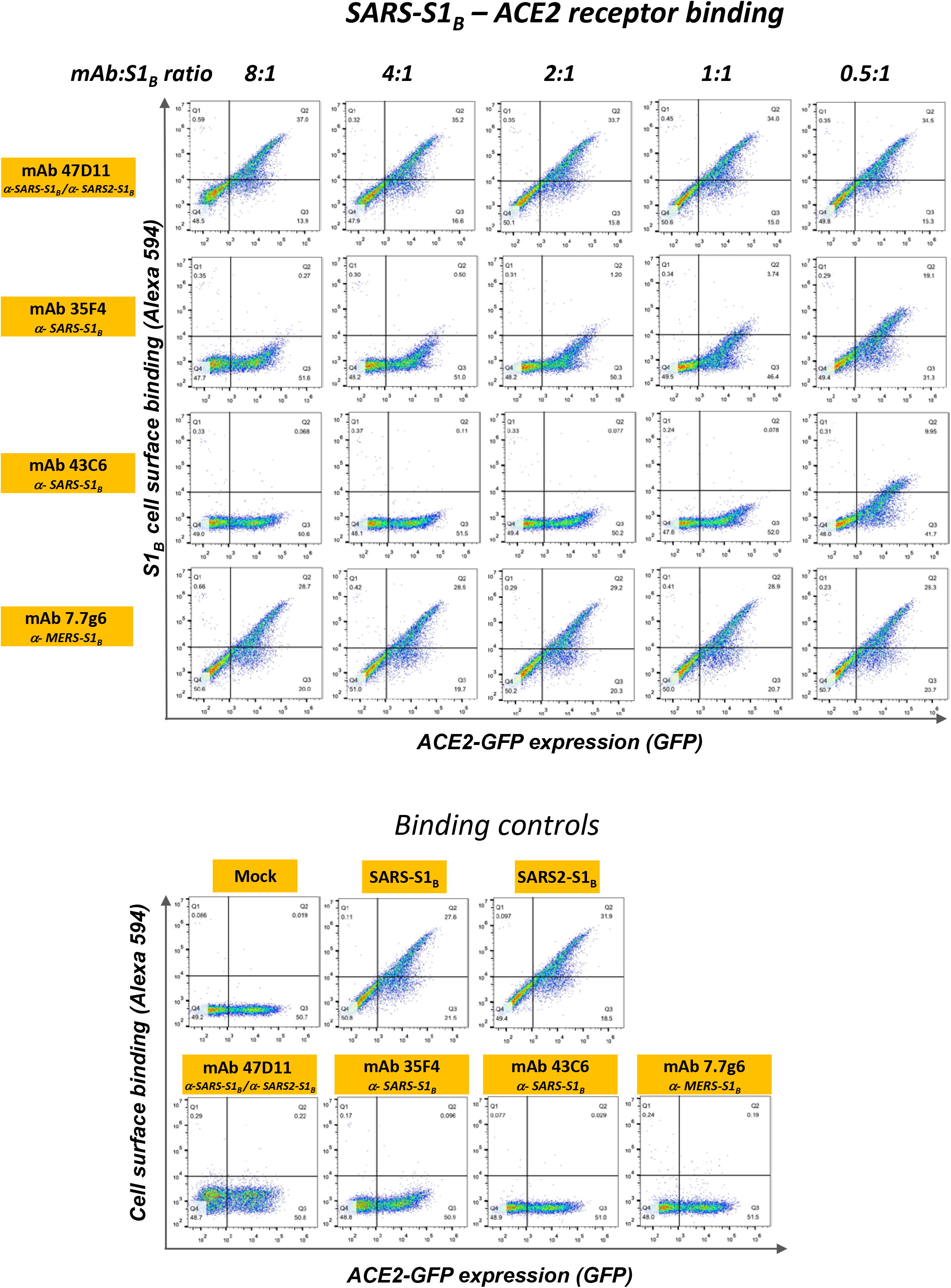
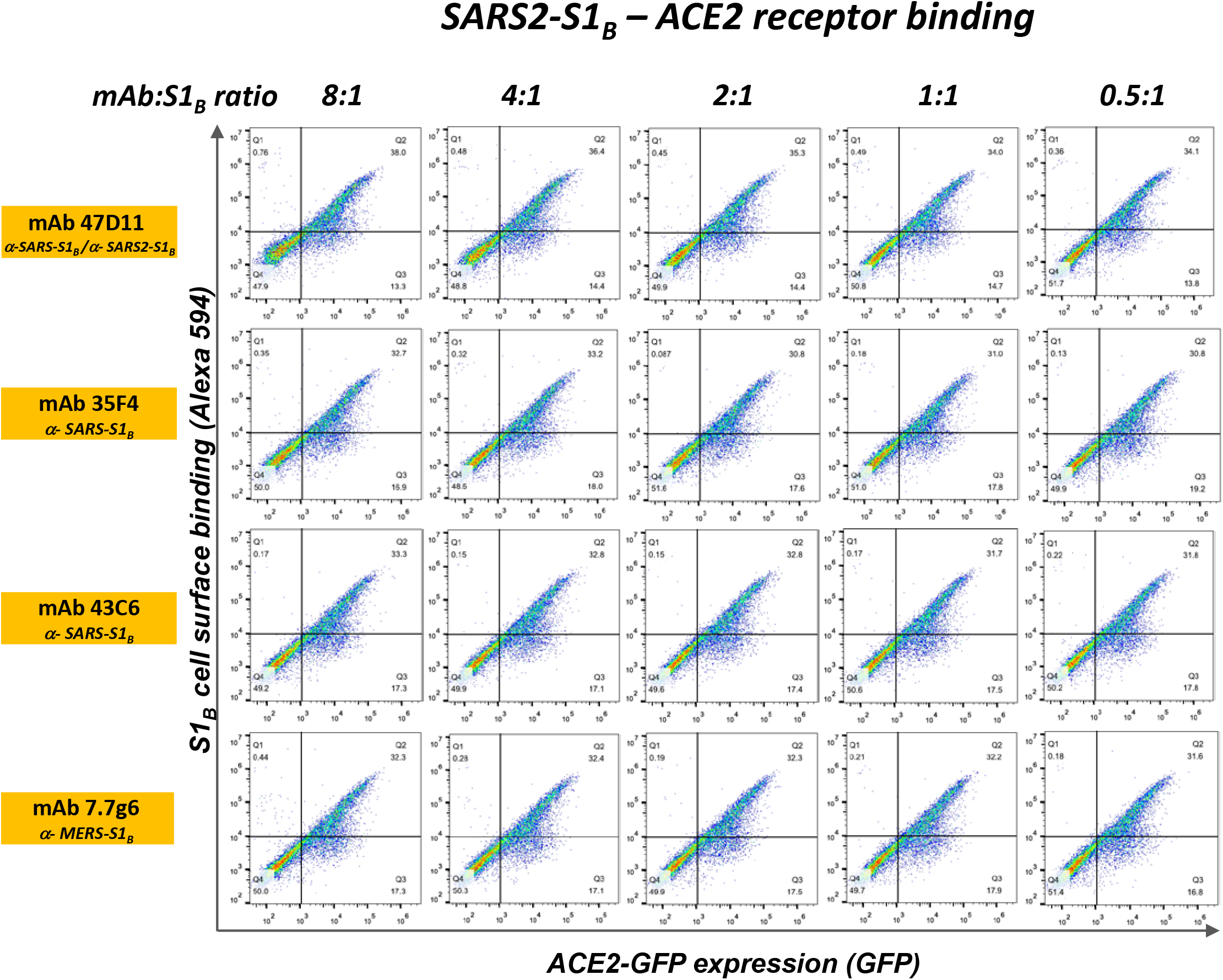
47D11 does not prevent binding of SARS-S1_B_ and SARS2-S1_B_ to ACE2-expressing cells. Human HEK-293T cells expressing human ACE2-GFP proteins (see Methods) were detached and fixed with 2% PFA, incubated with a fixed amount of human Fc-tagged S1_B_ domain of SARS-S or SARS2-S that was preincubated for 1h with mAb (mAbs 47D11, 35F4, 43C6, 7.7G6, in H2L2 format) at the indicated mAb:S1_B_ molar ratios, and analysed by flow cytometry using a Alexa Flour 594-conjugated secondary antibody targeting the human Fc tag. Cells are analysed for GFP expression (x-axis, GFP signal) and antibody binding (y-axis, Alexa 594 signal). Percentages of cells that scored negative, single positive, or double positive are shown in each quadrant. Binding controls include PBS-treated cells (mock), treatment of cells with SARS-S1_B_ and SARS2-S1_B_ in the absence of antibody, and cells treated with antibodies only. The experiment was performed twice, data from a representative experiment are shown.

**Suppl. Fig.5.**
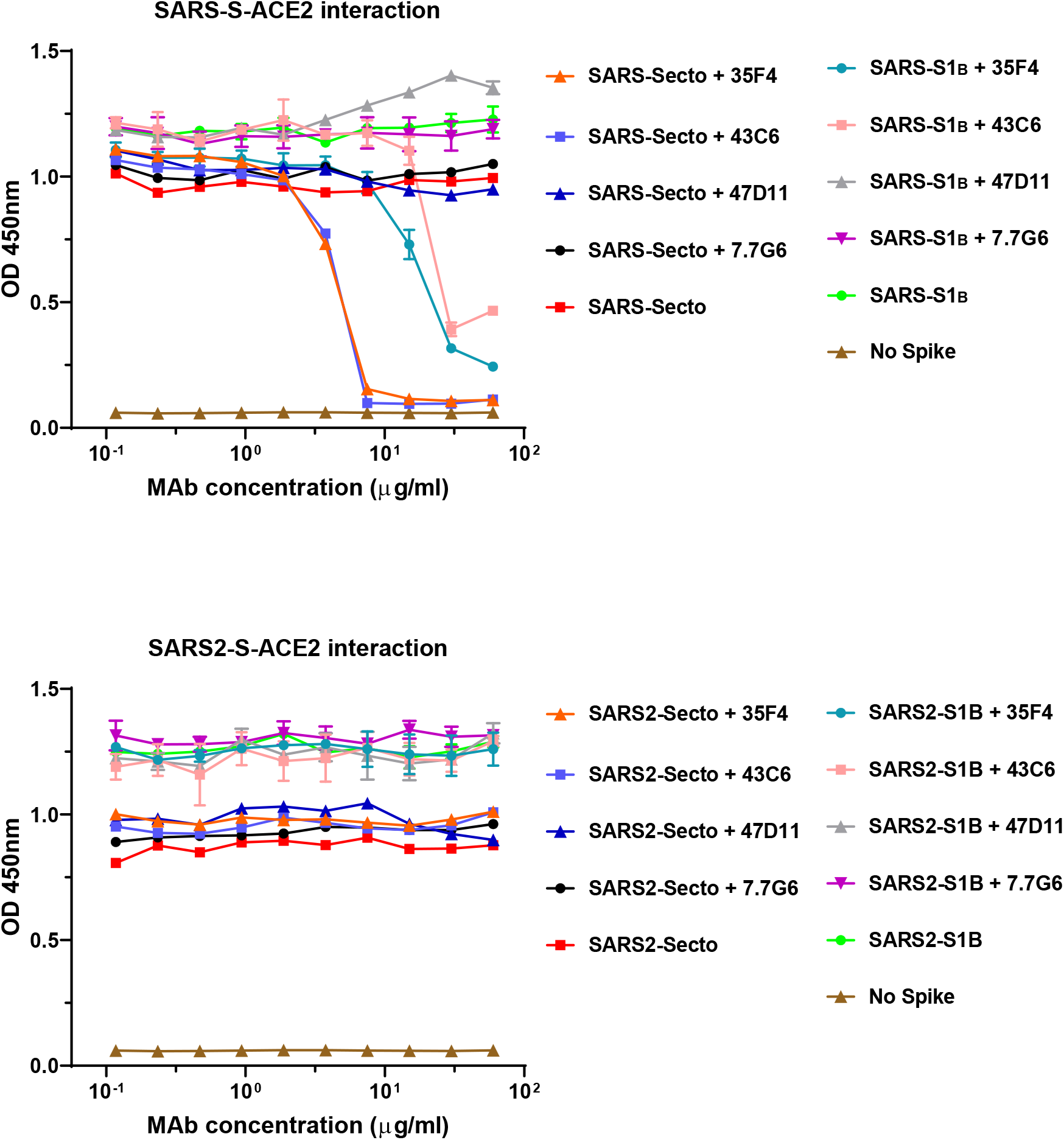
ELISA-based receptor binding inhibition assay. The ELISA-based receptor binding inhibition assay was performed as described previously with some adaptions^1^. Recombinant soluble human ACE2 was coated on NUNC Maxisorp plates (Thermo Scientific) at 4°C overnight. Plates were washed three times with PBS containing 0.05% Tween-20 and blocked with 3% BSA in PBS containing 0.1% Tween-20 at room temperature for 2 hours. Recombinant S_ecto_ and S1_B_ of SARS-S or SARS2-S (300 ng) and serially diluted mAbs (mAbs 47D11, 35F4, 43C6, 7.7G6, in H2L2 format) were mixed for 1 hour at RT, added to the plate for 1 t room temperature, after which the plates were washed three times. Binding to ACE2 was detected using HRP-conjugated StrepMAb (IBA) that recognizes the C-terminal Streptag on the S_ecto_ and S1_B_ proteins.

**Suppl. Fig.6.**
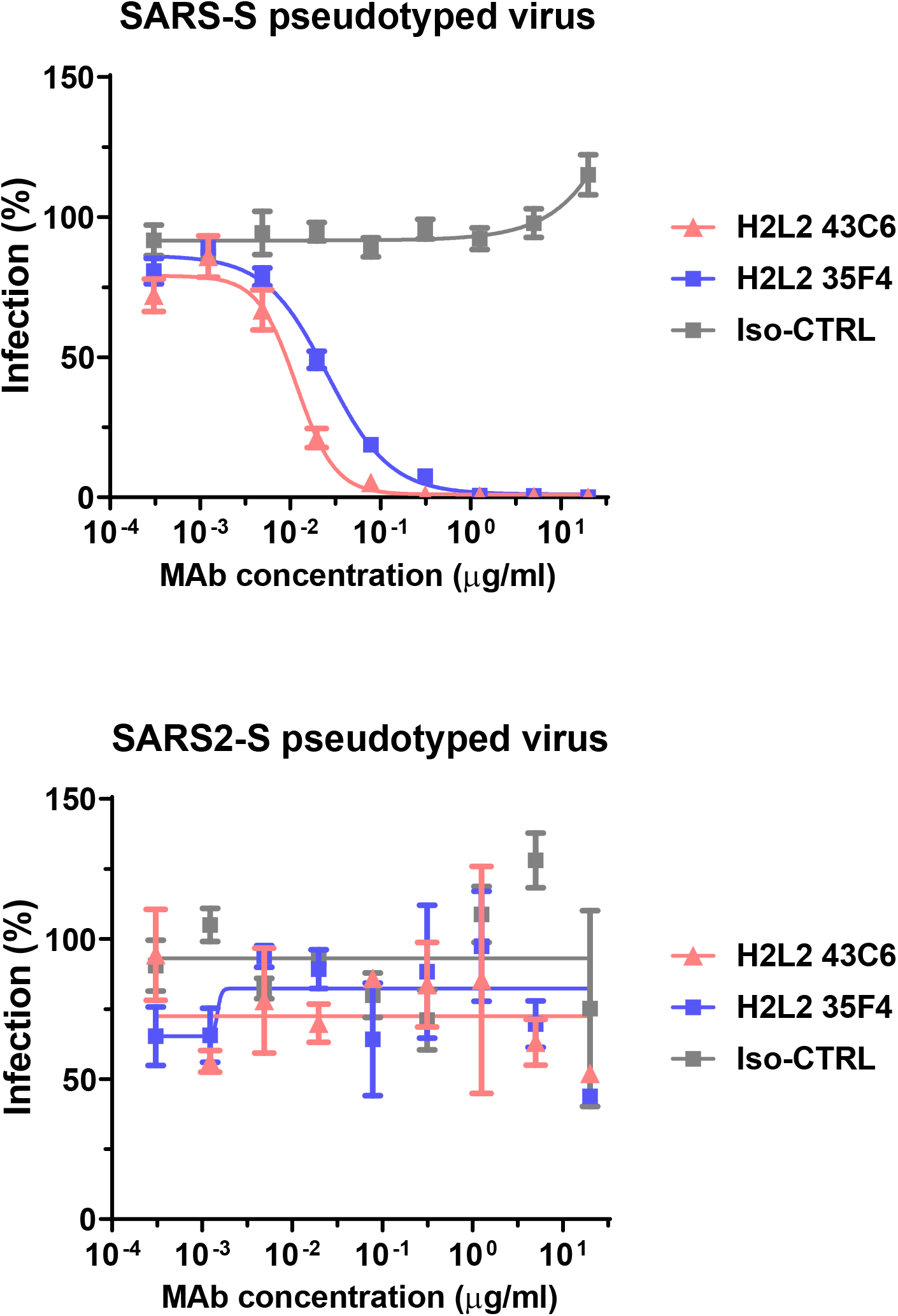
H2L2 monoclonal antibodies 35F4 and 43C6 neutralize SARS-CoV but not SARS-CoV-2. Antibody-mediated neutralization of infection of VSV particles pseudotyped with spike proteins of SARS-CoV (upper panel) and SARS-CoV-2 (lower panel) by the 35F4 and 43C6 H2L2 antibodies targeting SARS-S1 but not SARS2-S1 (see Suppl.Fig.1). An irrelevant antibody was taken along as a human IgG1 isotype control. Means ± SD of triplicates are shown.

**Suppl. Fig.7.**
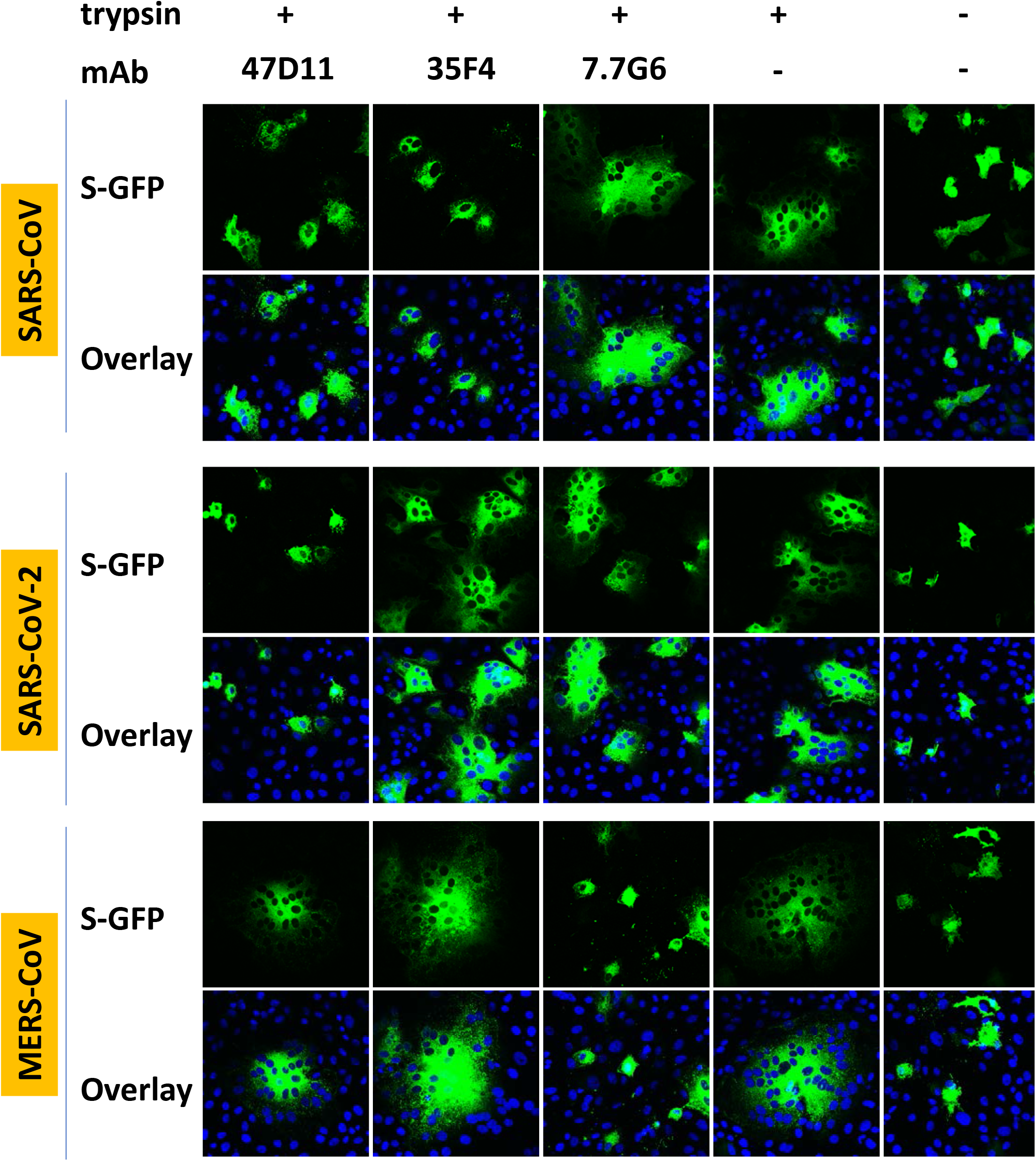
Cell-cell fusion inhibition assay. The cell-cell-fusion inbition assay was performed as described previously with some adaptations^1^. VeroE6 cells were seeded with density of 10^5^ cells per ml. After reaching 70~80% confluency, cells were transfected with plasmids encoding full length SARS-S, SARS2-S and MERS-S – C-terminally fused to GFP - using Lipofectamine 2000 (Invitrogen). The furin recognition site in the SARS2-S was mutated (R^682^RAR to A^682^AAR) to inhibit cleavage of the protein by endogenous furin and allow trypsin-induced syncytia formation. Two days post transfection, cells were pretreated DMEM only or DMEM with 20 μg/ml mAbs for 1 h and subsequently treated with DMEM with 15 μg/ml trypsin (to activate the spike fusion function) in the absence or presence of 20 μg/ml mAbs (47D11 crossreactive SARS-S and SARS2-S, 35F4 reactive to SARS-S, 7.7G6 reactive to MERS-S). After incubation at 37°C for 2 hrs, the cells were fixed with 2% PFA in PBS for 20 min at room temperature and stained for nuclei with 4,6-diamidino-2-phenylindole (DAPI). Cells expressing the S-GFP proteins were detected by fluorescence microscopy and S-mediated cell-cell fusion was observed by the formation of fluorescent) multi-nucleated syncytia. The fluorescence images were recorded using a Leica SpeII confocal microscope. The experiment was performed twice, data from a representative experiment are shown.

**Suppl. Fig.8.**
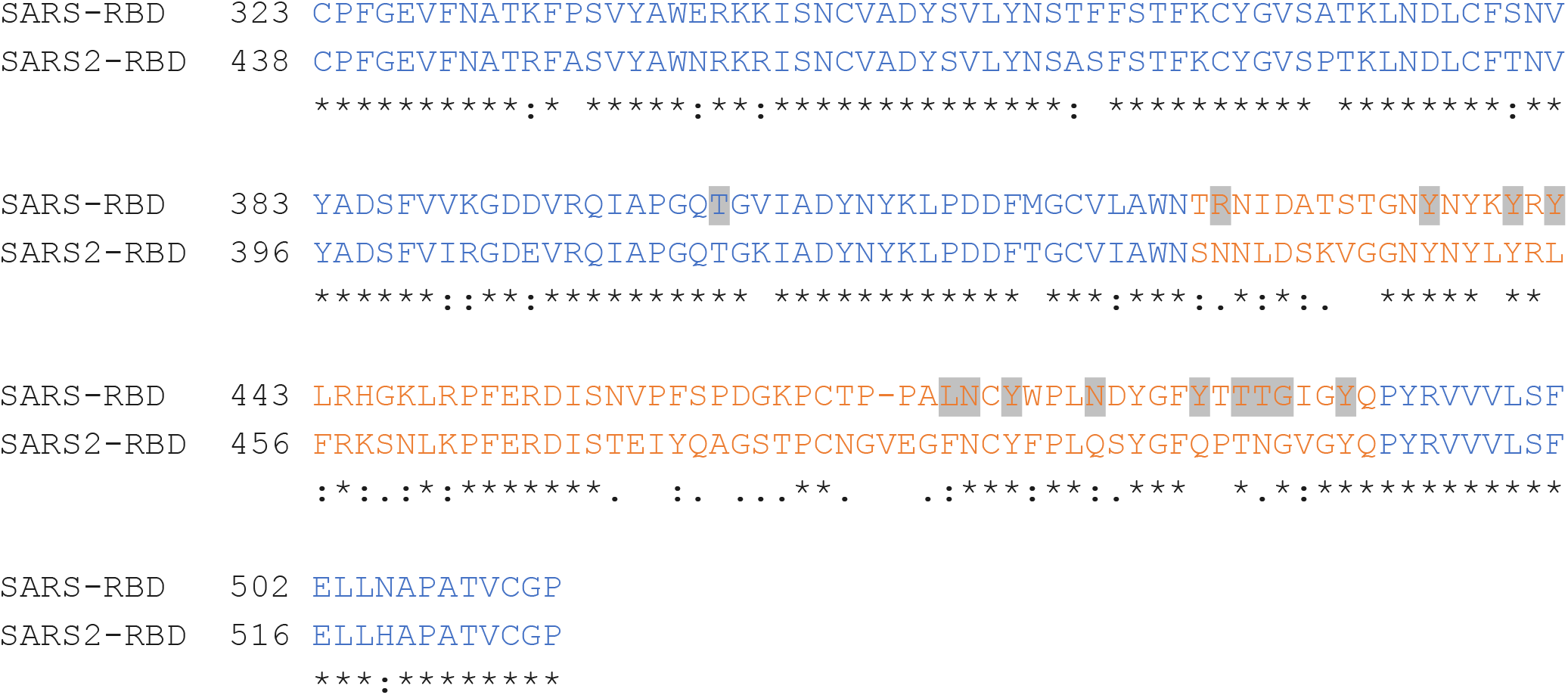
Protein sequence alignment of the S1_B_ receptor binding in (RBD) of the SARS-CoV and SARS-CoV-2 spike proteins by ClustalW. Numbering denotes the residue position in the full-length spike protein of SARS-CoV (Genbank: AAP13441.1) and SARS-CoV-2 (Genbank: QHD43416.1). Asterisks (*) indicated fully conserved residues, the colon symbol (:) indicates conservation between groups of very similar properties, and the period symbol (.) indicates conservation between groups of weakly similar properties. Sequences corresponding to the S1_B_ receptor binding core domain and the receptor binding subdomain are colored in blue and orange, respectively. The fourteen residues that are involved in binding of SARS-CoV S1_B_ human ACE2 are highlighted in grey^1^.

